# Enteropathogen antibody dynamics and force of infection among children in low-resource settings

**DOI:** 10.1101/522920

**Authors:** Benjamin F. Arnold, Diana L. Martin, Jane Juma, Harran Mkocha, John B. Ochieng, Gretchen M. Cooley, Richard Omore, E. Brook Goodhew, Jamae F. Morris, Veronica Costantini, Jan Vinjé, Patrick J. Lammie, Jeffrey W. Priest

## Abstract

Little is known about enteropathogen seroepidemiology among children in low-resource settings. We measured serological IgG responses to eight enteropathogens (*Giardia intestinalis, Cryptosporidium parvum, Entamoeba histolytica, Salmonella enterica*, enterotoxigenic *Escherichia coli, Vibrio cholerae, Campylobacter jejuni*, norovirus) in cohorts from Haiti, Kenya, and Tanzania. We studied antibody dynamics and force of infection across pathogens and cohorts. Enteropathogens shared common seroepidemiologic features that enabled between-pathogen comparisons of transmission. Overall, exposure was intense: for most pathogens the window of primary infection was <3 years old; for highest transmission pathogens primary infection occurred within the first year. Longitudinal profiles revealed significant IgG boosting and waning above seropositivity cutoffs, underscoring the value of longitudinal designs to estimate force of infection. Seroprevalence and force of infection were rank-preserving across pathogens, illustrating the measures provide similar information about transmission heterogeneity. Our findings suggest multiplex antibody assays are a promising approach to measure population-level enteropathogen transmission in serologic surveillance.

## Introduction

A broad set of viral, bacterial, and parasitic enteropathogens are leading causes of the global infectious disease burden, with the highest burden among young children living in lower income countries [1]. Infections that result in acute diarrhea and related child deaths drive disease burden estimates attributed to enteropathogens, but asymptomatic infections are extremely common and the full scope of sequelae is only partially understood [2,3]. Much of what we know about enteropathogen transmission is based on passive clinical surveillance, which reflects a small fraction of all infections. For example, antibody-based incidence of infection to *Salmonella enterica* and *Campylobacter jejuni* were 2-6 orders of magnitude higher than standard case-based surveillance in European populations [4–7], and a study of *Salmonella enterica* serotype Typhi in Fiji found similarly high discordance between antibody-based incidence and case-based surveillance [8]. A more complete picture of enteropathogen infection in populations would help understand drivers of transmission, disease burden, naturally acquired immunoprotection, as well as to design public health prevention measures, and measure intervention effects.

Stool-based, high-throughput PCR assays have helped solve the logistical difficulties of single-pathogen testing for enterics and have provided new insights into pathogen-specific infections and disease burden [2,3]. Yet, stool is not routinely collected in population-based surveys, and infection with many globally important enteric pathogens can be sufficiently rare and relatively short-lived to require designs with almost continuous surveillance [3,9]. At the same time, large-scale serological surveillance platforms create new opportunities for expanded enteropathogen surveillance alongside other infectious diseases [10,11]. These challenges and opportunities have generated interest in antibody-based measurement as a complement to PCR for population-based enteropathogen surveillance [12–15], and for endpoints in observational and randomized studies [16–22].

Many enteropathogens elicit a transiently elevated antibody response after infection that wanes over time. In lower transmission settings where antibody responses could be monitored longitudinally after distinct infections, *Salmonella enterica, Campylobacter jejuni, Cryptosporidium parvum,* and *Giardia intestinalis* (syn. *Giardia lamblia, Giardia duodenalis*) immunoglobulin G (IgG) levels in blood have been shown to wane over a period of months since infection; IgM and IgA levels decline even more quickly [6,23–27]. Compared with permanently immunizing infections such as measles, transient immunity adds a layer of complexity to seroepidemiologic inference and methods. To our knowledge, there has been no detailed study of enteropathogen seroepidemiology among children in low-resource settings where transmission is intense beginning early in life [3]. Such studies are needed to determine if serology is a viable approach to measure enteropathogen transmission in low-resource settings.

We conducted a series of analyses in cohorts from Haiti, Tanzania, and Kenya that measured serological antibody responses to eight enteropathogens using multiplex bead assays. Our objectives were to identify common patterns in antibody dynamics shared across enteropathogens and populations, and to evaluate serological methods to compare between-pathogen heterogeneity in infection, including estimates of force of infection. Our results provide new insights into the seroepidemiology of enteropathogens among children living in low-resource settings, and contribute advances to inform the design and analysis of surveillance efforts whose goal is to quantify heterogeneity in enteropathogen transmission through antibody response.

## Results

### Study populations

The analysis included measurements from cohorts in Haiti, Kenya, and Tanzania. Blood specimens were tested for IgG levels to eight enteropathogens using a multiplex bead assay on the Luminex platform (Table 1). The Haitian cohort included repeated measurements among children enrolled in a study of lymphatic filariasis transmission in Leogane from 1990–1999 [28,29]. Leogane is a coastal agricultural community west of Port au Prince. At the time of the study its population was approximately 15,000, most homes had no electricity and none had running water. In total, the Haiti study tested 771 finger prick blood specimens collected from 142 children ages birth to 11 years old, with each measurement typically separated by one year (median measurements per child: 5; range: 2 to 9). In Kenya, a 2013 prospective trial of locally-produced, in-home ceramic water filters enrolled 240 children in a serological substudy [30]. Study participants were identified through the Asembo Health and Demographic Surveillance System, which is located in a rural part of Siaya County, western Kenya along the shore of Lake Victoria. Only 29% of the population had piped drinking water (public taps), water source contamination with *E. coli* prevailed (93% of samples tested), and the average age children began consuming water was 4 months [30]. Children aged 4 to 10 months provided blood specimens at enrollment, and again 6 to 7 months later (n=199 children measured longitudinally). In Tanzania, 96 independent clusters across 8 trachoma-endemic villages in the Kongwa region were enrolled in a randomized trial to study the effects of annual azithromycin distribution on *Chlamydia trachomatis* infection [31]. The population is very rural, and water is scarce in the region: at enrollment, 69% of participants reported their primary drinking water source, typically an unprotected spring, was >30 minutes’ walk one-way. From 2012 to 2015, the Tanzania study tested dried blood spots from between 902 and 1,577 children ages 1–9 years old in annual cross-sectional surveys (total measurements: 4,989). Although children could have been measured repeatedly over the four-year study in Tanzania, they were not tracked longitudinally. There was no evidence that the Kenya and Tanzania interventions reduced enteropathogen antibody response (Supplementary Information File 1), so this analysis pooled measurements from the study arms in each population.

**Table 1.**
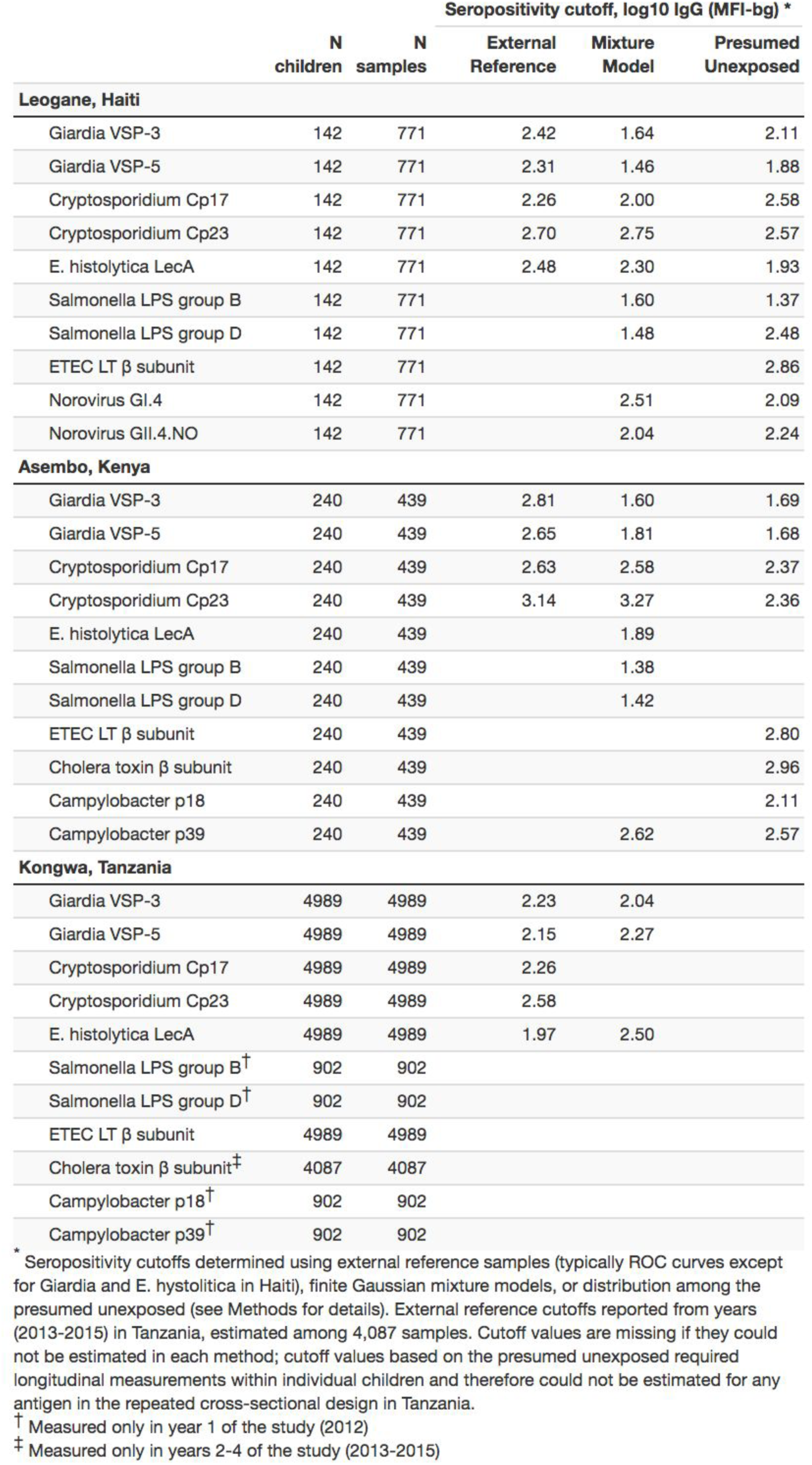
Number of children and samples tested, and estimated seropositivity cutoffs by country and IgG antigen included in the seroepidemiologic analyses.

### Age-dependent shifts in population antibody distributions

We estimated seropositivity cutoffs using three approaches: receiver operator characteristic (ROC) curve analyses for *Giardia, Cryptosporidium,* and *Entamoeba histolytica* including a panel of external, known positive and negative specimens, as previously reported [14,30]; Gaussian mixture models [32] fit to measurements among children ages 0–1 year old to ensure a sufficient number of unexposed children; and, presumed seronegative distributions among children who experienced large increases in antibody levels (Table 1). Classification agreement was high between the different approaches (agreement >95% for most comparisons; Supplementary Information File 2).

Among children <2 years old, antibody levels clearly distinguished seronegative and seropositive subpopulations, but there were not distinct seronegative and seropositive subpopulations by age 3 years for most pathogens measured in Haiti (Figure 1) and Tanzania (Figure 1 – supplement 1). By age 3 years, the majority of children were seropositive to *Cryptosporidium*, enterotoxigenic *Escherichia coli* heat labile toxin β subunit (ETEC LT β subunit), and norovirus GI.4 and GII.4; in all cases antibody distributions were shifted above seropositivity thresholds. In contrast, there was a qualitative change in the antibody response distributions to *Giardia, E. histolytica, Salmonella* and *Campylobacter* with increasing age, shifting from a bimodal distribution of seronegative and seropositive groups among children ≤1 year old to a unimodal distribution by age 3 years and older (Figure 1, Figure 1 – supplement 1). A direct comparison of age-dependent shifts in antibody distributions to *Giardia* VSP-3 antigen and *Chlamydia* pgp3 antigen in Tanzania illustrates stark differences in enteropathogen-generated immune responses versus pathogens like *Chlamydia* that elicit a response that consistently differentiates exposed and unexposed subpopulations as children age (Figure 1 – supplement 2). Distributions of IgG levels in the younger Kenyan cohort (ages 4–17 months) showed distinct groups of seropositive and seronegative measurements for most antigens (Figure 1 – supplement 3). IgG responses to ETEC LT β subunit and cholera toxin β subunit were near the maximum of the dynamic range of the assay for nearly all children measured in the three cohorts (Figure 1, Figure 1 – supplement 1, Figure 1 – supplement 3), and these IgG levels waned as children aged, presumably from adaptive immunity (Figure 1, Figure 1 – supplement 1).

**Figure 1.**
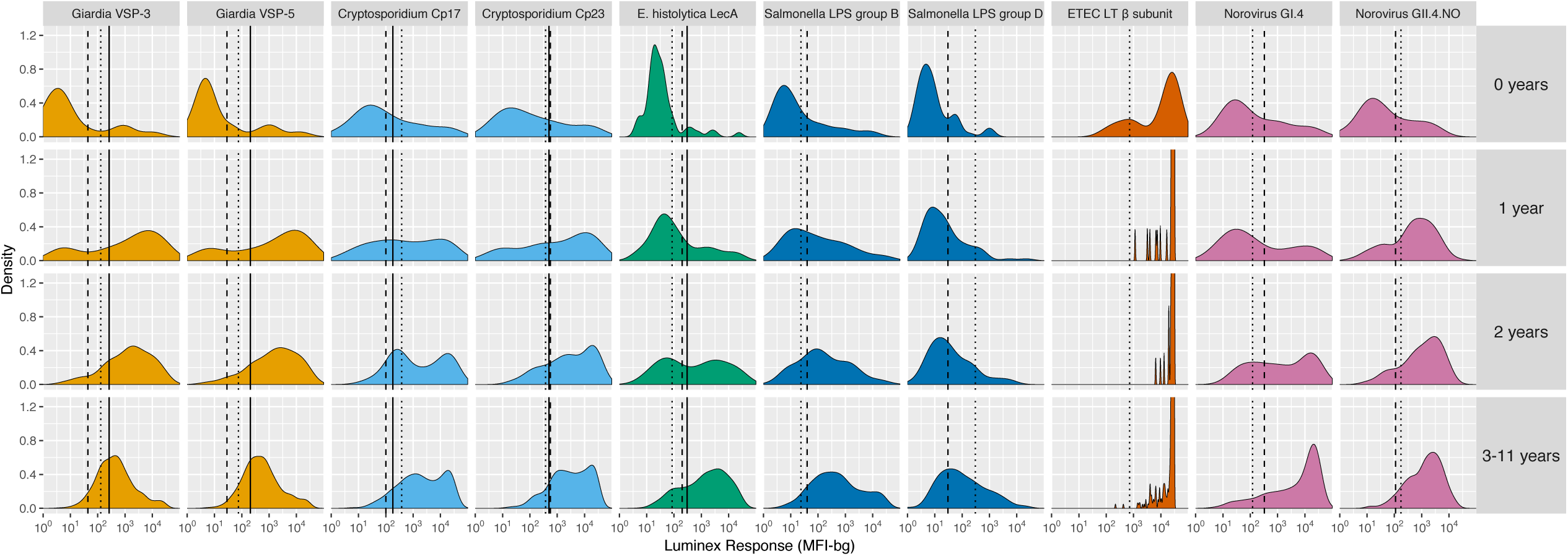
Age-stratified, IgG distributions among a longitudinal cohort of 142 children ages birth to 11 years in Leogane, Haiti, 1990– 1999. IgG response measured in multiplex using median fluorescence units minus background (MFI-bg) on the Luminex platform in 771 specimens. Vertical lines mark seropositivity cutoffs are based on ROC analyses (solid), finite Gaussian mixture models (heavy dash), or distribution among presumed unexposed (light dash). Mixture models failed to converge for ETEC heat labile toxin *β* subunit. Created with notebook (https://osf.io/dk54y) and data (https://osf.io/3nv98). Supplement 1 shows similar distributions from the Tanzania study. Supplement 2 contrasts *Giardia* VSP-3 distributions with *C. trachomatis* pgp3 distributions. Supplement 3 shows distributions from the Kenyan cohort.

### Joint variation in antibody response

We hypothesized that IgG responses to closely related antigens would co-vary but that IgG responses to unrelated antigens would be uncorrelated. Joint variation in individual-level IgG responses aligned with hypothesized relationships based on antigenic overlap and shared epitopes. Responses to *Giardia* VSP antigens were strongly correlated in Haiti (Spearman rank: *ρ*=0.99), Kenya (*ρ*=0.84) and Tanzania (*ρ*=0.97), as would be expected for antigens with conserved conformational epitopes (Figure 2). *Cryptosporidium* (Cp17, Cp23) and *Campylobacter* (p18, p39) antigens were strongly correlated, but high within-individual variability suggests that measuring responses to multiple unique recombinant protein antigens yields more information about infection than measuring responses to one alone (Figure 2). High correlation between *Salmonella* LPS Groups B and D, between norovirus GI.4 and GII.4, and between ETEC and *V. cholerae* likely reflected antibody cross-reactivity. Correlation could also result from multiple previous infections with different *Salmonella* serogroups or different norovirus genogroups. A comparison across all antigens revealed no other combinations with high correlation (Supplementary Information File 3).

**Figure 2.**
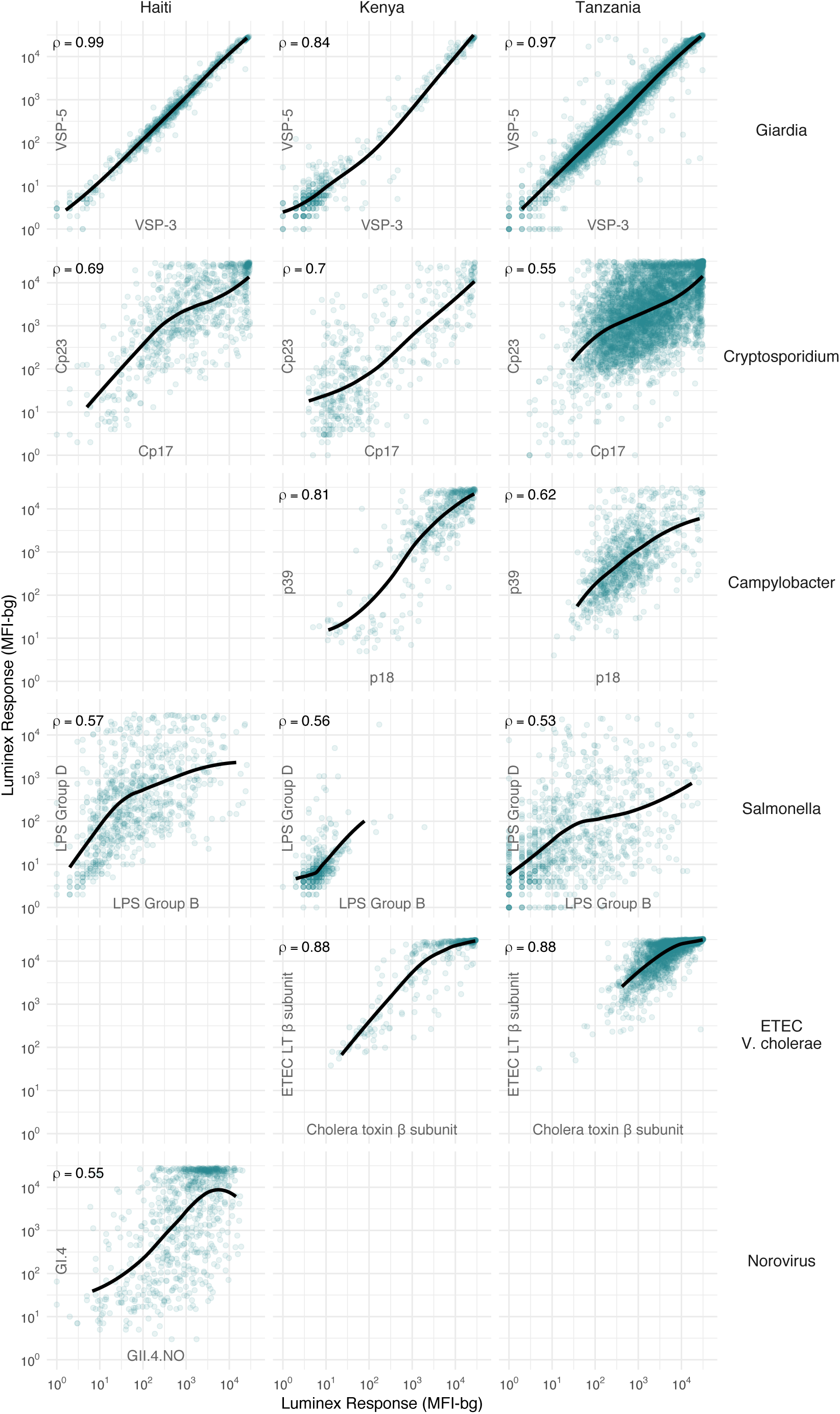
Joint distributions of select enteric pathogen antibody responses among children in three cohorts from Haiti, Kenya, and Tanzania. Each panel includes Spearman rank correlations (*ρ*) and locally weighted regression smoothers with default parameters, trimmed to 95% of the data to avoid edge effects. Antibody response measured in multiplex using median fluorescence units minus background (MFI-bg) on the Lu-minex platform. Empty panels indicate that the antibodies were not measured in that cohort. Supplementary Information File 3 includes all pairwise comparisons. Created with notebook (https://osf.io/hv9ce) and data (https://osf.io/3nv98, https://osf.io/2q7zg, https://osf.io/kv4d3).

We excluded cholera β toxin antibody responses from remaining analyses because of the difficulty of interpreting its epidemiologic measures in light of high levels of cross-reactivity with ETEC LT β subunit. Heat labile toxin-producing ETEC is very common among children in low-resource settings [3], and there was no documented transmission of cholera in the study populations during measurement periods.

### Birth to three years of age: a key window of antibody acquisition and primary seroconversion

For most pathogens, mean IgG levels and seroprevalence rose quickly and plateaued by ages 1 to 3 years in Haiti (Figure 3), Kenya (Figure 3 – supplement 1), and Tanzania (Figure 3 – supplement 2). Despite enormous individual-level variation, age-dependent mean IgG curves exhibited characteristic shapes seen across diverse pathogens, and reflected high levels of early-life exposure [15]. In Haiti, seroprevalence ranged from 66% (*E. histolytica*) to 100% (ETEC LT β subunit) by age 3 years (Figure 3B), and in Tanzania, the majority of 1 year olds were already seropositive for *Giardia* (77%) and *Cryptosporidium* (85%) (Figure 3B – supplement 2). There was some evidence of maternally-derived IgG among children under 6 months old with a drop in mean IgG levels by age, but this pattern was only evident for norovirus GI.4 in Haiti (Figure 3A) and *Cryptosporidium* Cp17 and Cp23 in Kenya (Figure 3A – supplement 1). Age-dependent mean IgG responses to many pathogens declined from a young age, presumably from exposure early in life and acquired immunity. Mean IgG levels declined after age 1 year for *Giardia* in Haiti (Figure 3A), *Giardia* and *Campylobacter* in Tanzania (Figure 3A – supplement 2).

**Figure 3.**
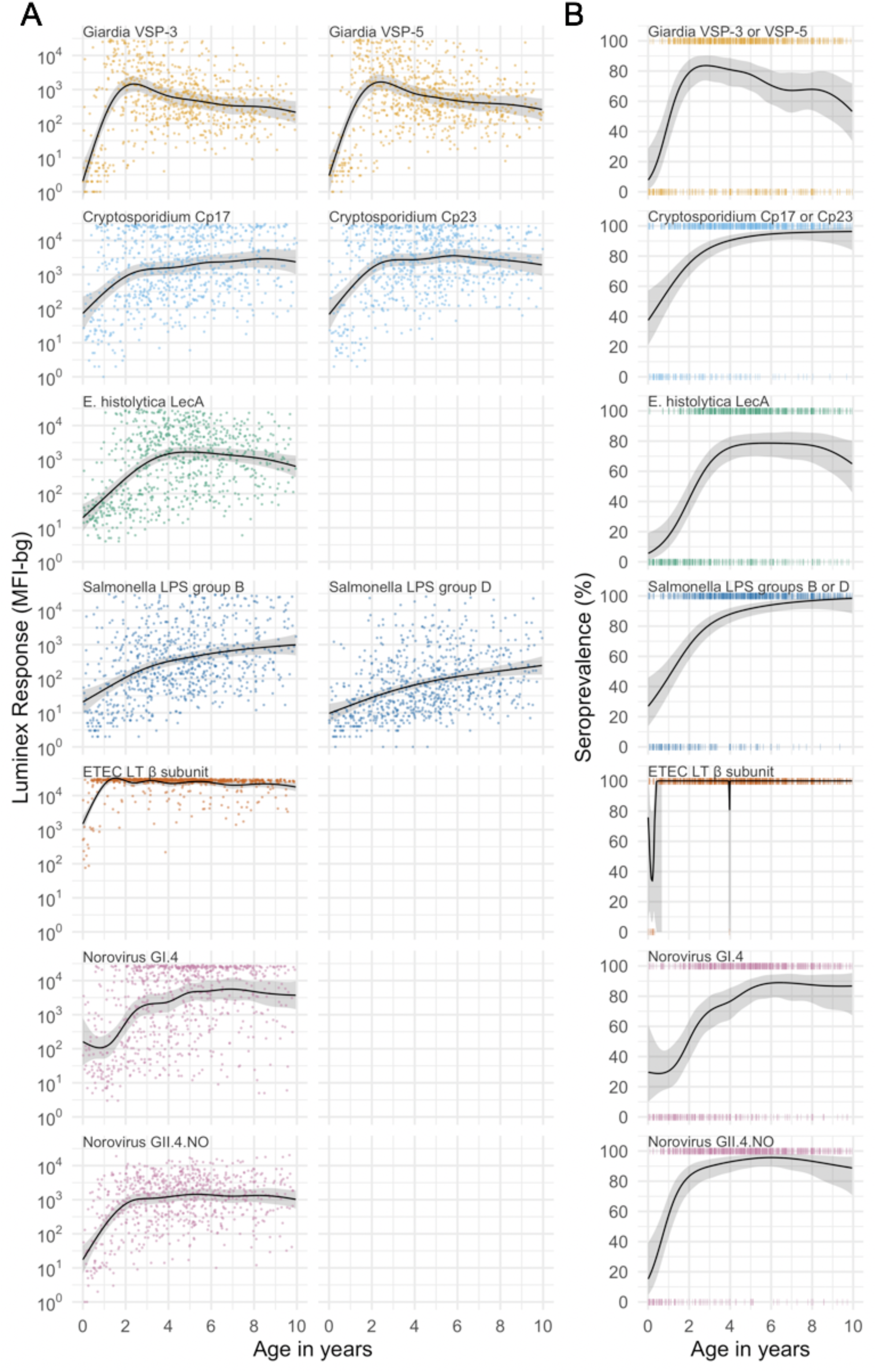
Age dependent curves for geometric means (**A**) and seroprevalence (**B**), estimated with cubic splines among children ages birth to 10 years in Leogane, Haiti 1990-1999. Shaded bands are approximate, simultaneous 95% credible intervals. IgG response measured in multiplex using median fluorescence units minus background (MFI-bg) on the Luminex platform (N=771 measurements from 142 children). Created with notebook (https://osf.io/6c7j8) and data (https://osf.io/3nv98). Data for some antigens measured among children <5 years previously published [8]. Supplement 1 includes similar curves from Kenya; Supplement 2 includes similar curves from Tanzania.

### Longitudinal antibody dynamics show significant boosting and waning above seropositivity cutoffs

Based on age-dependent shifts in IgG distributions (Figure 1), we hypothesized that conversion of IgG levels to seropositive and seronegative status could mask important dynamics of enteropathogen immune response above seropositivity cutoffs, particularly among ages beyond the window of primary infection. In Haiti and Kenya we examined longitudinal IgG profiles among children. In the Haitian cohort, which was followed beyond the window of primary infection, children commonly had >4-fold increases and decreases in IgG while remaining above seropositivity cutoffs—a pattern observed across pathogens but particularly clear for *Cryptosporidium* (Figure 4). In Kenya, 4-fold increases in IgG largely coincided with a change in status from seronegative to seropositive, presumably because increases in IgG followed primary infection in the young cohort ages 4–17 months (Figure 4 – supplement 1). Many Kenyan children exhibited >4-fold increases and decreases in IgG response to *Campylobacter* p18 and p39 antigens above the seropositivity cutoff, a result of earlier primary infection and/or additional infection and boosting during the study period (Figure 4 – supplement 1).

**Figure 4.**
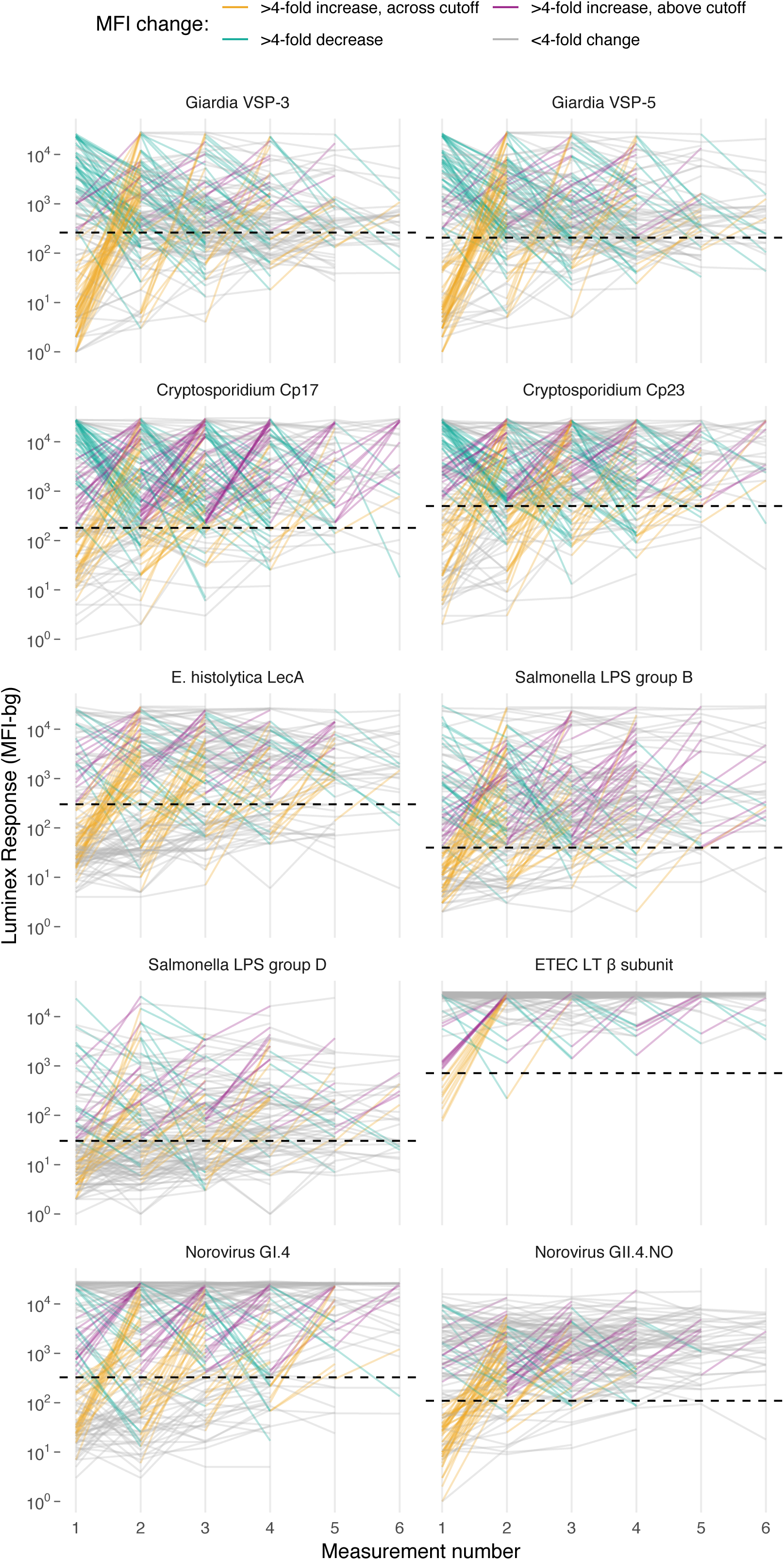
Longitudinal changes in IgG response over six repeated measurements among 142 children ages 0-11 years in Leogane, Haiti, 1990-1999. Measurements were spaced by approximately 1 year (median spacing = 1, IQR = 0.7, 1.3). Horizontal dashed lines mark seropositivity cutoffs for each antibody. The number of children measured at each visit was: n_1_=142, n_2_=142, n_3_=140, n_4_=131, n_5_=111, n_6_=66); 29 children had >6 measurements that are not shown. IgG response measured in multiplex using median fluorescence units minus background (MFI-bg) on the Luminex platform. Created with notebook (https://osf.io/mf7g2), which includes additional visualizations, and data (https://osf.io/3nv98). Supplement 1 includes individual antibody trajectories in the Kenya cohort.

### Serological estimates of force of infection

The seroconversion rate, an instantaneous rate of seroconversion among those who are susceptible, is one estimate of a pathogen’s force of infection and a fundamental epidemiologic measure of transmission [33]. Serologically derived force of infection is useful for pathogens that commonly present asymptomatically, such as many enteric infections. Across diverse pathogens, steeper age-seroprevalence curves typically reflect higher transmission intensity [34,35], and overall seroprevalence equals the area under the age-seroprevalence curve (a summary measure) [15]. We therefore hypothesized that seroprevalence and prospectively estimated force of infection should embed similar information about infection heterogeneity across pathogens. We also hypothesized that standard methods to estimate force of infection from age-structured seroprevalence would underestimate force of infection derived from longitudinal data because of significant antibody boosting and waning above seropositivity cutoffs.

Longitudinal designs in Haiti and Kenya enabled us to use individual child antibody profiles to estimate average rates of prospective seroconversion and seroreversion during the studies. We defined incident seroconversions and seroreversions as a change in IgG across a pathogen’s seropositivity cutoff and estimated force of infection as incident changes in serostatus divided by person-time at risk. In a secondary analysis, we defined incident boosting as a ≥4-fold increase in IgG to a final level above a seropositivity cutoff and incident waning as ≥4-fold decrease in IgG from an initial level above a seropositivity cutoff. The secondary definition captured large changes in IgG above seropositivity cutoffs, which aligned with repeated boosting and waning observed in the Haitian cohort (Figure 4).

We found a rank-preserving relationship between pathogen seroprevalence and average force of infection in Kenya and Haiti (Figure 5). Overall levels and steepness of the relationship differed between cohorts, presumably because Kenya measurements were within a window of primary infection for most children (4–17 months) whereas Haiti measurements extended from birth to 11 years and captured lower incidence periods with overall higher seroprevalence as children aged. Force of infection varied widely across pathogens in Kenya, ranging from 0.1 seroconversions per year for *E. histolytica* to >5 for *Campylobacter* (Figure 5).

**Figure 5.**
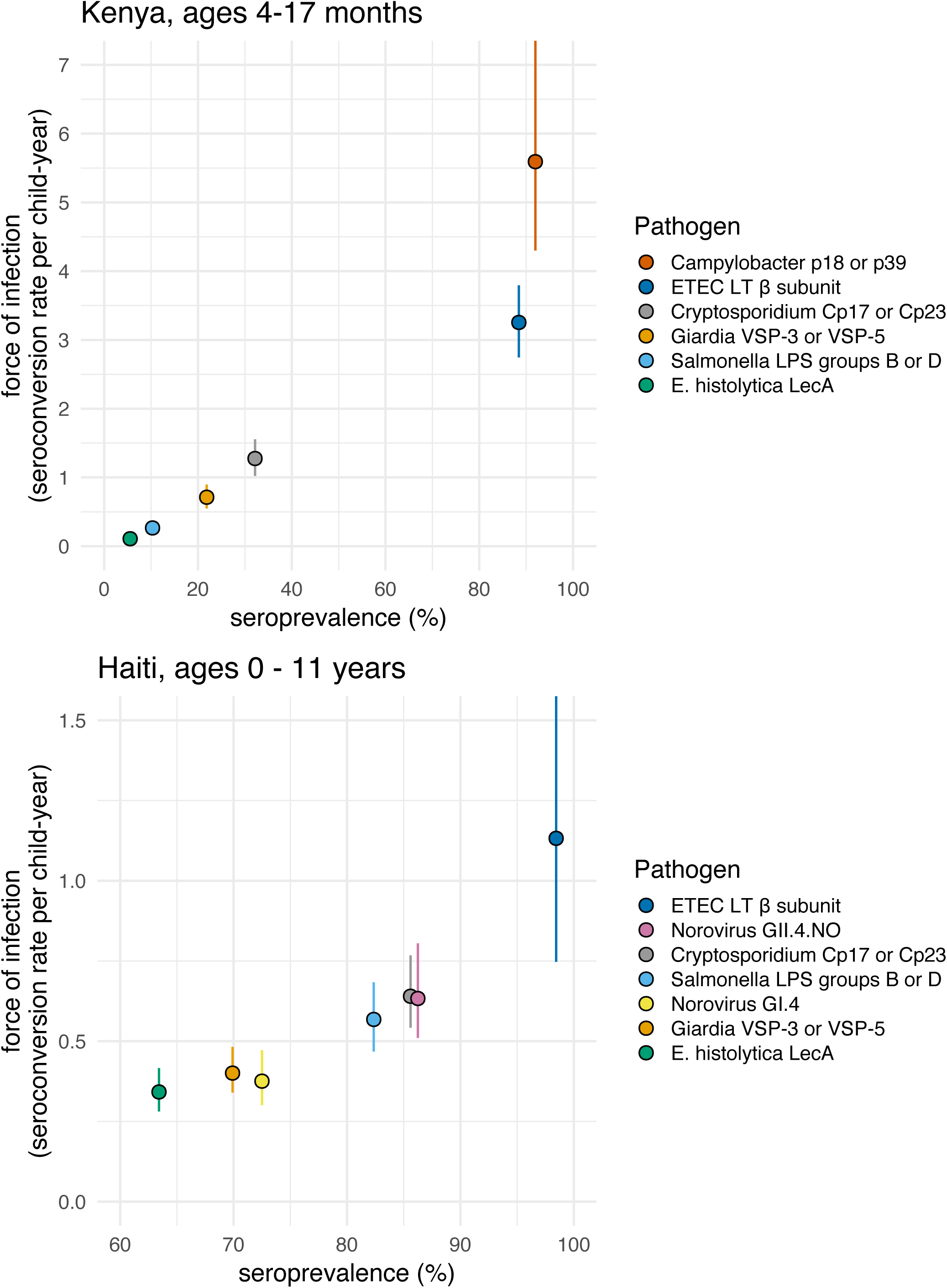
Average force of infection versus seroprevalence for enteropathogens measured in the Kenya and Haiti cohorts. Force of infection estimated from prospective seroconversion rates. Dashed lines show linear fits across pathogens. Note that axis scales differ between panels to best-illustrate the estimates. Created with notebook (https://osf.io/jp9kf) and data (https://osf.io/2q7zg, https://osf.io/3nv98).

In Haiti, force of infection ranged from 0.3 *E. histolytica* seroconversions per year to 1.1 ETEC seroconversions per year (Figure 5). Force of infection estimated from 4-fold changes in IgG led to more events and higher rates compared with those estimated from seroconversion alone (Table 2). For example, *Cryptosporidium* incident cases increased from 70 to 190 (a 2.7 fold increase) and the average rate increased from 0.6 (95% CI: 0.5, 0.8) to 0.8 (95% CI: 0.7, 0.9) per child-year (a 1.3 fold increase) when using a 4-fold IgG change criteria because of substantial IgG boosting and waning above the seropositivity cutoff (Figure 4).

**Table 2.** Seroincidence rates per child-year among children ages 0-11 years in Leogane, Haiti, 1990-1999. Rates estimated using two definitions of incident changes in serostatus, one based on seroconversion and a second based on 4-fold change in IgG levels. Ratio of incident cases and seroincidence rates divides the second definition by the first.

| Pathogen | Seropositivity cutoff * |  |  | 4 Fold Change in IgG levels † |  |  | Ratio of cases | Ratio of rates |
| --- | --- | --- | --- | --- | --- | --- | --- | --- |
|  | Child-years | Incident cases | Rate (95% CI) | Child-years | Incident cases | Rate (95% CI) |  |  |
| Seroconversion / boosting |  |  |  |  |  |  |  |  |
| Giardia VSP-3 or VSP-5 | 269.6 | 108 | 0.40 (0.34, 0.48) | 275.9 | 118 | 0.43 (0.34, 0.54) | 1.1 | 1.1 |
| Cryptosporidium Cp17 or Cp23 | 109.3 | 70 | 0.64 (0.54, 0.77) | 236.6 | 190 | 0.80 (0.70, 0.92) | 2.7 | 1.3 |
| E. histolytica LecA | 283.7 | 97 | 0.34 (0.28, 0.42) | 299.9 | 120 | 0.40 (0.32, 0.50) | 1.2 | 1.2 |
| Salmonella LPS groups B or D | 132.1 | 75 | 0.57 (0.47, 0.68) | 224.4 | 140 | 0.62 (0.51, 0.75) | 1.9 | 1.1 |
| ETEC LT β subunit | 9.7 | 11 | 1.13 (0.75, 1.82) | 32.1 | 33 | 1.03 (0.72, 1.53) | 3.0 | 0.9 |
| Norovirus GI.4 | 213.0 | 80 | 0.38 (0.30, 0.47) | 257.0 | 118 | 0.46 (0.36, 0.57) | 1.5 | 1.2 |
| Norovirus GII.4.NO | 105.8 | 67 | 0.63 (0.51, 0.80) | 148.5 | 109 | 0.73 (0.58, 0.94) | 1.6 | 1.2 |
| Seroreversion / waning |  |  |  |  |  |  |  |  |
| Giardia VSP-3 or VSP-5 | 441.6 | 91 | 0.21 (0.17, 0.25) | 291.9 | 126 | 0.43 (0.35, 0.54) | 1.4 | 2.1 |
| Cryptosporidium Cp17 or Cp23 | 586.1 | 29 | 0.05 (0.03, 0.07) | 276.9 | 166 | 0.60 (0.51, 0.70) | 5.7 | 12.1 |
| E. histolytica LecA | 395.2 | 43 | 0.11 (0.08, 0.15) | 307.8 | 70 | 0.23 (0.17, 0.29) | 1.6 | 2.1 |
| Salmonella LPS groups B or D | 544.3 | 25 | 0.05 (0.03, 0.07) | 349.1 | 86 | 0.25 (0.19, 0.30) | 3.4 | 5.4 |
| ETEC LT β subunit | 702.1 | 2 | 0.00 (0.00, 0.01) | 649.7 | 22 | 0.03 (0.02, 0.05) | 11.0 | 11.9 |
| Norovirus GI.4 | 464.9 | 28 | 0.06 (0.03, 0.09) | 359.9 | 58 | 0.16 (0.11, 0.22) | 2.1 | 2.7 |
| Norovirus GII.4.NO | 574.3 | 19 | 0.03 (0.02, 0.05) | 476.9 | 40 | 0.08 (0.06, 0.11) | 2.1 | 2.5 |

\* Incident changes in serostatus defined by crossing seropositivity cutoffs.
† Incident changes in serostatus defined by a 4-fold increase or decrease in IgG levels (MFI-bg), with incident boosting episodes restricted to changes that ended above the seropositivity cutoff and incident waning episodes restricted to changes that started from above the seropositivity cutoff.

We evaluated whether model-based force of infection estimates from age-structured seroprevalence could accurately recover estimates from the longitudinal analyses. We focused on the Kenya cohort since children were measured repeatedly during the ages of primary infection and because longitudinal force of infection and seroreversion rate estimates varied considerably across pathogens (Figure 6A). We estimated force of infection from seroprevalence curves using methods developed for cross-sectional, “current status” data, a common approach in serosurveillance of vaccine preventable diseases [33], malaria [34], and dengue [36]. Force of infection estimates from semiparametric spline models were similar to estimates from the longitudinal analysis for all pathogens, but had substantially wider confidence intervals owing to the loss of information from ignoring the longitudinal data structure (Figure 6B). Parametric approaches including an exponential survival model [37] and a reversible catalytic model [34] yielded narrower confidence intervals than the semiparametric model but tended to underestimate force of infection compared with longitudinal estimates (Figure 6B). Across pathogens, model-based force of infection estimates derived from seroprevalence were rank-preserving compared with nonparametric longitudinal analyses.

**Figure 6.**
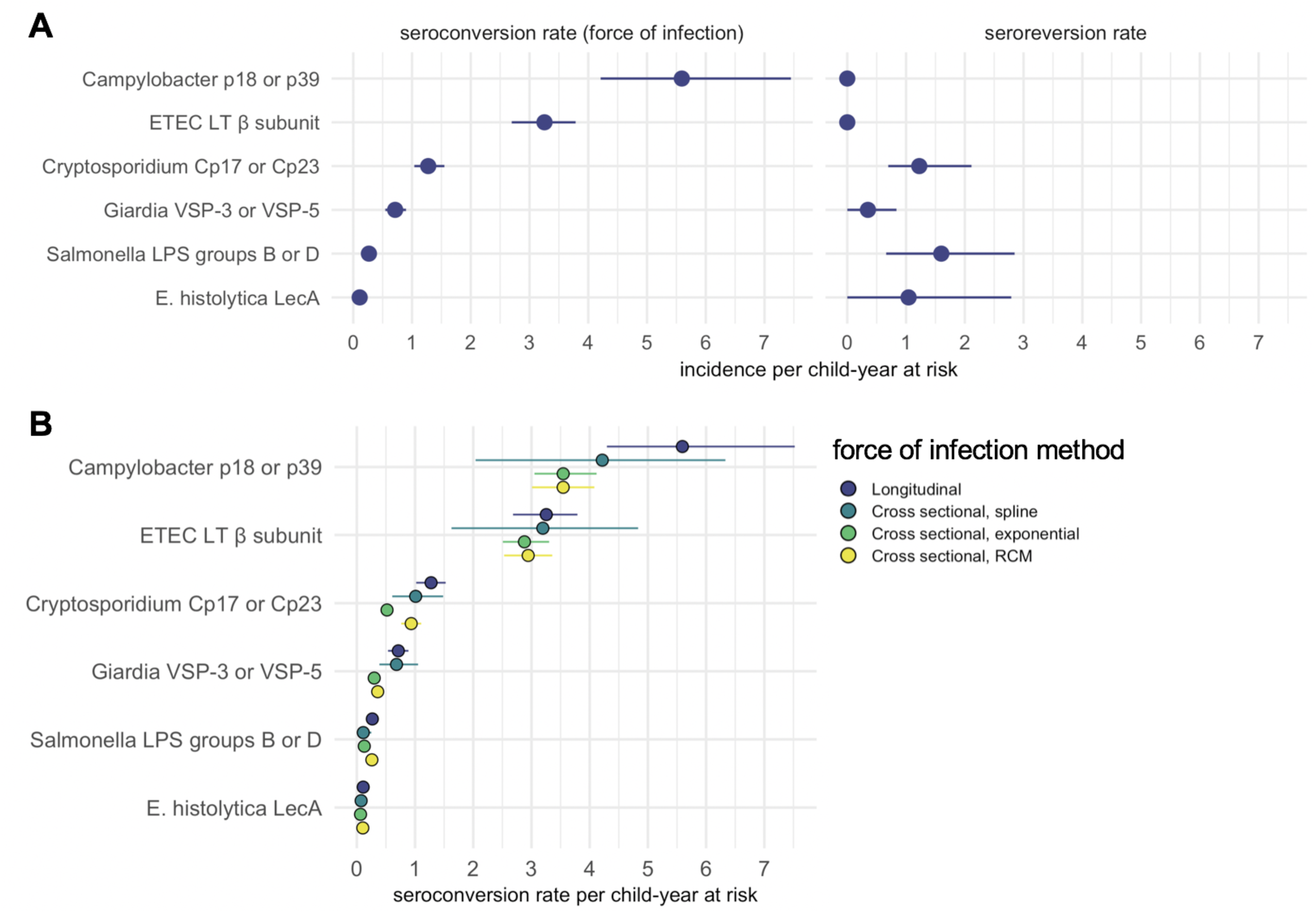
**A** Enteropathogen seroconversion and seroreversion rates among 199 children ages 4 to 17 months measured longitudinally in Asembo, Kenya, 2013. The seroconversion rate is a measure of a pathogen’s force of infection. **B** Comparison of force of infection estimation methods that included longitudinal analysis (as in panel A), plus cross-sectional estimators derived from age-specific seroprevalence curves using semiparametric cubic splines (spline), a parametric constant rate survival model (exponential), and a reversible catalytic model (RCM) that assumed a constant seroconversion rate and fixed seroreversion rate (estimated from the prospective analysis). Lines mark 95% confidence intervals. IgG response measured in multiplex using median fluorescence units minus background (MFI-bg) on the Luminex platform (N=398 measurements from 199 children). Created with notebooks (https://osf.io/sqvj7, https://osf.io/y5ukv) and data (https://osf.io/2q7zg).

## Discussion

In cohorts from Haiti, Kenya, and Tanzania we identified consistent patterns in IgG responses that provide new insights into enteropathogen seroepidemiology among children in low-resource settings. Most population-level heterogeneity in IgG levels and seroconversion was between birth and 3 years, reflecting high transmission and early life primary infection. For particularly high transmission pathogens (e.g., ETEC, *Campylobacter*), most variation in IgG levels was observed among children <1 year old. In these study populations, endemic force of infection for enteropathogens was as high, and in many cases several fold higher, than force of infection estimated during epidemics of new dengue serotype introductions in Nicaragua and Peru (approximately 0.4 to 0.6 seroconversions per year) [36,38]. The shift of IgG distributions from bimodal to unimodal for many pathogens (*Giardia, Cryptosporidium, E. histolytica,* and *Campylobacter*), resulting from a combination of antibody boosting, waning and adaptive immunity, complicates the interpretation of seroprevalence at older ages: among older children a seronegative response could either mean the children were never exposed or they were previously exposed but antibody levels waned below seropositivity cutoffs. Significant boosting and waning of antibody levels above seropositivity cutoffs identified through longitudinal profiles in Haiti and Kenya reinforce the value of longitudinal designs to derive antibody distributions among unexposed and to estimate force of infection. Among children in low-resource settings, seroprevalence and seroconversion rates are thus most clearly interpretable when estimated within a window of primary infection, and the window likely closes by age 2–3 years.

Seroepidemiologic measures that can be estimated from cross-sectional surveys are of particular interest for infectious diseases because most large-scale, population-based serosurveillance platforms use cross-sectional designs [11]. Our results show that seroprevalence and force of infection estimated from seroprevalence models adequately summarize between-pathogen heterogeneity in transmission when compared with longitudinal estimates of force of infection. Pathogens with fastest rising mean IgG and seroprevalence with age (e.g., ETEC, *Campylobacter,* norovirus GII.4; Figure 3, Figure 3 – supplement 1) had highest force of infection measured prospectively over the study period, and seroprevalence was rank-preserving with prospective force of infection in both Haiti and Kenya (Figure 5). Seroprevalence alone thus appears to be sufficient to assess relative pathogen transmission if measured in an age range that captures ample heterogeneity in response (in these cohorts <3 years old). Our findings align with modeling studies of other infectious diseases such as malaria [34], trachoma [35], and dengue [36], and suggest that enteropathogens share similar seroepidemiologic features conducive to population-based surveillance in cross-sectional surveys despite different underlying immunology. Two caveats are that cross-sectional methods considered in this study will generally underestimate force of infection because they fail to rigorously account for seroreversion and incident boosting above seropositivity cutoffs (Figure 6B), and seroprevalence is not a clear marker of cumulative exposure as children age beyond the window of primary infection because of waning IgG responses.

The relative intensity of pathogen transmission based on serological measures in this analysis aligns with relative differences in infection from large-scale molecular testing in stool. For example, rapid rises in IgG levels, seroprevalence, and force of infection for ETEC and *Campylobacter* beginning very early in life is consistent with their prominence in molecular testing of stool specimens in the MAL-ED studies [3,39]. Higher force of infection for norovirus GII.4 compared with norovirus GI.4 in Haiti is consistent with the relative prominence of circulating genogroups [40,41]. The results lend additional support to the idea that broader serological testing in existing serosurveillance platforms represents a useful opportunity to measure enteropathogen transmission in populations without routine stool-based testing [10,11].

These analyses had limitations. High correlation between *Salmonella* LPS Groups B and D, between ETEC LT β subunit and cholera toxin β subunit, and between norovirus GI and norovirus GII at the individual level (Figure 2) shows that for these pathogens seroepidemiologic analyses will be less specific than analyses based on molecular detection. *Salmonella* LPS Groups B and D have antigenic overlap in their lipopolysaccharides [42], and ETEC LT β subunit and cholera toxin β subunit are known to be immunologically cross-reactive [43]. There was no known cholera transmission in the study populations, so we assumed the elevated responses to the cholera toxin β subunit reflected exposure to LT-producing ETEC; this assumption could only be confirmed with specific measures of cholera infection. Norovirus GI and GII virus-like particles are antigenically different, but cross-reactivity between norovirus genogroups is possible [44]. In addition, a recent study from Uganda found high seroprevalence for both GI and GII norovirus suggesting repeated infections by viruses of the same or different genogroup could potentially boost cross-reactive antibody production [45]. Despite lower specificity between genogroups or serogroups in these cases, the consistency of antibody relationships with predictions based on antigenic overlap, and absence of correlation between unrelated antibodies (Supplementary Information File 3), lends support to the internal validity of the assays.

Another limitation is that without measures of patent infection in stool we were unable to compare multiple independent measures of transmission. The Haiti and Tanzania cohorts were not originally designed to assess enteropathogens, and antibody measurements were not paired with measures of clinical symptoms of diarrhea or with measures of patent infection in stool. In Kenya, stool-based pathogen testing was limited and low prevalence prevented detailed comparisons between serological and molecular results: for example, only 14 children were positive by PCR to *Cryptosporidium* [30]. All included antigens have been characterized extensively with respect to patent infections among adults and children in other settings (details in Materials and Methods). Furthermore, we observed high levels of consistency across cohorts in seroepidemiologic patterns, and consistency with general age-dependent patterns documented across diverse pathogens for which transmission begins early in life [15,33,34,36]. Together, these observations suggest that the antibody dynamics and force of infection estimates from IgG responses reflect actual pathogen transmission in these cohorts, but paired stool and blood testing would provide a more definitive test.

High-resolution, longitudinal assessment of paired enteropathogen infection and antibody measurements among children could provide valuable, additional insights into the pathogen-specific antibody dynamics following primary- and secondary infections. Analyses of Plasmodium falciparum [46,47] and dengue virus [48] illustrate how paired, longitudinal measurements of patent infection and antibody response enable richer characterizations of antibody dynamics and pathogen transmission. Studies that pair stool-based molecular testing with antibody response could also assess whether approaches to estimate enteropathogen incidence from cross-sectional samples that rely on estimates of antibody decay with time since infection [6,49] could be used among children in low-resource settings. A similar approach was recently described to estimate cholera incidence among all ages [50], but broader development across enteropathogens and among young children remains an open area of research.

## Conclusions

Among children in Haiti, Kenya, and Tanzania, antibody-based measures of enteropathogen infection reflected high transmission with primary exposure to most pathogens occurring by age 1–2 years. In low-resource populations, seroincidence rates and force of infection estimated beyond the age range of primary infection ideally should account for IgG boosting and waning above seropositivity cutoffs. Antibodies are a promising approach to measure population-level enteropathogen infection, and seroepidemiologic measures of heterogeneity and transmission are central considerations for their use in trials or in serologic surveillance. Our findings show that for most enteropathogens studied, the ideal window to measure heterogeneity in antibody response closes by age two years in low-resource settings, and studies that plan to estimate force of infection should favor longitudinal designs with multiple measurements in this early age window.

## Materials and Methods

### Ethics statement

In Haiti, the human subjects protocol was reviewed and approved by the Ethical Committee of St. Croix Hospital (Leogane, Haiti) and the institutional review board at the US Centers for Disease Control and Prevention (CDC). After listening to an overview of the study, individuals were asked for verbal consent to participate. Verbal consent was deemed appropriate by both review boards because of low literacy rates in the study population. With each longitudinal visit, the study team re-consented participants before specimen collection. Mothers provided consent for children under 7, and children 7 years and older provided additional verbal assent. In Kenya, the human subjects protocol was reviewed and approved by institutional review boards at the Kenya Medical Research Institute (KEMRI) and at the US CDC. Primary caretakers provided written informed consent for their infant child’s participation in the trial and blood specimen collection and testing [30]. The original trial was registered at clinicaltrials.org (NCT01695304). In Tanzania, the human subjects protocol was reviewed and approved by the Institute for Medical Research Ethical Review Committee in Dar es Salaam, Tanzania and the institutional review board at the US CDC. Parents of enrolled children provided consent, and children 7 years and older also provided verbal assent before specimen collection.

### Multiplex bead assays

#### Antigens

All antigens used have been well-characterized in previous studies. Lipopolysaccharides (LPS) from Group D Salmonella *enterica* serotype Enteritidis [51], LPS from Group B *S. enterica* serotype Typhimurium [51], and recombinant heat labile toxin β subunit protein [15,52,53] from enterotoxigenic *Escherichia coli* (ETEC LT β subunit) were purchased from Sigma Chemical (St. Louis, MO). Recombinant *Giardia* VSP-3 and VSP-5 [54,55] and *Cryptosporidium* Cp17 and Cp23 antigens [56–58] were expressed and purified as previously described [25,59]. Recombinant *Campylobacter* p18 and p39 antigens [60–62] were expressed and purified as previously described [17]. *E. histolytica* LecA antigen [14,63,64] was kindly provided by William Petri (University of Virginia) and Joel Herbein (TechLab). Virus-like particles from norovirus GI.4 and GII.4 New Orleans were purified from a recombinant baculovirus expression system [15,65,66].

#### Haiti

Sera from the Haiti cohort study were diluted 1:400 and analyzed by multiplex bead assay as described in detail elsewhere [14,15,29]. *Salmonella* LPS and ETEC LT β subunit antigens were coupled to SeroMap (Luminex Corp, Austin, TX) beads in buffer containing 0.85% NaCl and 10 mM Na_2_HPO_4_ at pH 7.2 (PBS) using 120 micrograms for 1.25 × 10^7^ beads using the methods described by Moss and colleagues [67]. Coupling conditions and externally defined cutoff values for the *Giardia, Cryptosporidium*, and *E. histolytica* antigens as well as for the *Schistosoma japonicum* glutathione-*S*-transferase (GST) negative control protein have been previously reported [14].

#### Kenya

For the Kenya study, an optimized bead coupling technique using less total protein was performed in buffer containing 0.85% NaCl and 25 mM 2-(N-morpholino)-ethanesulfonic acid at pH 5.0. The β subunit protein from cholera toxin was purchased from Sigma Chemical. The GST negative control protein (15 μg), *Cryptosporidium* Cp17 (6.8 μg) and Cp23 (12.5 μg) proteins and the *Campylobacter* p39 protein (25 μg) were coupled to 1.25 × 10^7^ beads using the indicated protein amounts [17]. The *Giardia, E. histolytica*, ETEC, cholera, and *Campylobacter* p18 proteins were coupled using 30 μg of protein per 1.25 × 10^7^ beads. *Salmonella* LPS B and LPS D were coupled to the same number of beads using 60 μg and 120 μg, respectively. Blood spot elutions from the Kenya study were diluted to a final serum concentration of 1:400 (assuming 50% hematocrit) and analyzed by multiplex bead assay as described by Morris and colleagues [30].

#### Tanzania

For the Tanzania study, the same conditions described in the Kenya study were used to couple antigens from *Giardia, Cryptosporidium, E. histolytica*, ETEC β toxin subunit, cholera β toxin subunit, GST, and *Salmonella* LPS group B and LPS group D. *Campylobacter* p39 and p18 were both coupled at 25 μg per 1.25 × 10^7^ beads in buffer containing 0.85% NaCl and 25 mM 2-(N-morpholino)-ethanesulfonic acid at pH 5.0. Dried blood spots were eluted in the casein-based buffer described previously [24] and samples were diluted to either 1:400 serum dilution with 50 µl run per well for year 1, or 1:320 serum dilution with 40 µl run per well for years 2-4. The incubation steps, washes, and data collection methods used in the multiplex bead assay were performed as described previously [24]. All samples were run in duplicate, and the average median fluorescence intensity minus background (MFI-bg) value was recorded. The Tanzania study used different bead lots in year 1 and years 2-4; we confirmed that the use of different bead lots had no influence on the results (Supplementary Information File 1).

### Antibody distributions and determination of seropositivity

We transformed IgG levels to the log_10_ scale because the distributions were highly skewed. Means of the log-transformed data represent geometric means. We summarized the distribution of log_10_ IgG response using kernel density smoothers. In the Tanzania and Haiti cohorts, where children were measured across a broad age range, we stratified IgG distributions by each year of age <3 years to examine age-dependent changes in the population distributions. To assess potential cross-reactivity between antigens, we estimated pairwise correlations between individual-level measurements in each cohort using a Spearman rank correlation [68] and visualized the relationship for each pairwise combination with locally weighted regression fits [69].

We compared three approaches to estimate seropositivity cutoffs. *Approach 1:* External known positive and negative specimens were used to determine seropositivity cutoffs for *Giardia* VSP-3 and VSP-5 antigens, *Cryptosporidium* Cp17 and Cp23 antigens, and *E. histolytica* LecA antigen. Cutoffs were determined using ROC analysis as previously described [14,30] for all antigens except for LecA, VSP-3, and VSP-5 in Haiti; in these cases, the mean plus 3 standard deviations of 65 specimens from citizens of the USA with no history of foreign travel were used to estimate cutoffs [14]. *Approach 2:* We fit a 2-component, finite Gaussian mixture model [32] to the antibody distributions among children 0-1 years old, and estimated seropositivity cutoffs using the lower component’s mean plus three standard deviations. The rationale for restricting the mixture model estimation in Haiti and Tanzania to children 0-1 years old was based on initial inspection of the age-stratified IgG distributions that revealed a shift from bimodal to unimodal distributions by age 3 (Figure 1). This approach ensured that there was a sufficiently large fraction of unexposed children in the sample to more clearly estimate a distribution among seronegative children. *Approach 3:* In the longitudinal Haiti and Kenya cohorts we identified children <1 year old who presumably seroconverted, defined as an increase in MFI-bg values of +2 or more on the log_10_ scale. A sensitivity analysis showed that an increase of 2 on the log_10_ scale was a conservative approach to identify seroconversion for most antibodies considered in this study; an increase of between 0.3 to 2.16 MFI-bg lead to optimal agreement with ROC-based and mixture model-based classifications in Kenya, and an increase of 0.92 to 2.41 led to optimal agreement across antigens and references in Haiti (Supplementary Information File 4). We then used the distribution of measurements before seroconversion to define the distribution of IgG values among the presumed unexposed. We used the mean log_10_ MFI-bg plus three standard deviations of the presumed unexposed distribution as a seropositivity cutoff. We summarized the proportion of observations that were in agreement between the three classification approaches, and estimated Cohen’s Kappa [70]. Additional details and estimates of seropositivity cutoff agreement are reported in Supplementary Information File 2. Mixture models failed to estimate realistic cutoff values if there was an insufficient number of unexposed children, which was the case for ETEC LT β subunit and cholera toxin β subunit in all cohorts, and for nearly all antigens in Tanzania where the study did not enroll children <1 year old (Table 1).

In analyses of seroprevalence and seroconversion, we classified measurements as seropositive using ROC-based cutoffs if available, and mixture model-based cutoffs otherwise. There were three exceptions. By age 1 year, a majority of children across the cohorts had IgG levels near the maximum of the assay’s dynamic range for enterotoxigenic *Escherichia coli* heat labile toxin β subunit (ETEC LT β toxin) and *Vibrio cholerae* toxin β subunit. The absence of a sufficient number of unexposed children to ETEC LT β toxin, *V. cholerae*, and in some cases *Campylobacter* led mixture models either to not converge or to estimate unrealistically high seropositivity cutoffs beyond the range of quantifiable levels. For these pathogens, we used seropositivity cutoffs estimated from presumed unexposed measurements in the longitudinal Haiti and Kenya cohorts (approach 3, above). High levels of agreement between classifications (Supplementary Information File 2) meant results were insensitive to choice of approach in these cohorts. We classified children as seropositive to *Giardia, Cryptosporidium, Campylobacter*, or *Salmonella* if antibody levels against either of the antigens from each pathogen were above estimated seropositivity cutoffs.

### Age-dependent antibody levels and seroprevalence curves

We estimated mean IgG levels and seroprevalence by age using semiparametric cubic splines in a generalized additive model, specifying binomial errors for seroprevalence, and random effects for children or clusters in the case of repeated observations [71,72]. We also estimated the relationships by age using a stacked ensemble approach called “super learner” that included a broader and more flexible library of machine learning algorithms [15,73,74], and found similar fits to cubic splines. We estimated approximate, simultaneous 95% credible intervals around the curves using a parametric bootstrap from posterior estimates of the model parameter covariance matrix [75]. Supplementary Information File 5 includes additional details.

### Force of infection from longitudinal data

In the Kenya and Haiti longitudinal cohorts, we estimated prospective seroconversion rates as a measure of force of infection by dividing the number of children who seroconverted by the person-time at risk between measurements. We defined incident seroconversions and seroreversions as a change in IgG across a pathogen’s seropositivity cutoff. Vaccine immunogenicity and pathogen challenge studies among healthy adults often use a 4-fold increase in antibody levels (difference of +0.6 on the log_10_ scale) as a criterion for seroconversion [76–78]. In a secondary analysis aimed to capture significant changes above a pathogen’s seropositivity cutoff, we defined incident boosting episodes as a ≥4-fold increase in IgG to a final level above a seropositivity cutoff, and incident waning episodes as ≥4-fold decrease in IgG from an initial level above a seropositivity cutoff. In the secondary definition, individuals were considered at risk for incident boosting episode if they were seronegative, if they experienced a ≥4-fold increase in IgG in their first measurement period, or if they experienced a ≥4-fold decrease in IgG in a preceding period (Haiti). To estimate person-time at risk used for rates and force of infection, we assumed incident changes were interval-censored and occurred at the midpoint between measurements. We estimated 95% confidence intervals for rates with 2.5 and 97.5 percentiles of a nonparametric bootstrap [79] distribution that resampled children with replacement to account for repeated observations.

### Force of infection from age-structured seroprevalence in Kenya

In the Kenya cohort, we estimated force of infection through age-structured seroprevalence using multiple approaches. There is a long history methods development to estimate force of infection from age-dependent seroprevalence [33], which is of particular interest to large-scale, cross-sectional surveillance platforms [11]. Our rationale was to determine if force of infection estimates from age-structured seroprevalence were comparable to estimates from the longitudinal analysis based on incident changes in serostatus.

As we show in Supplementary Information File 6, the age dependent seroprevalence curve is the difference between the cumulative distribution functions of seroconversion times and seroreversion times. In a special case of no seroreversion, age-specific seroprevalence is thus the cumulative hazard function. The age-specific force of infection can then be estimated as the hazard of seroconverting at age *A = a*: λ(*a*) = *F*′(*a*) / [1 – *F*(*a*)], where *F*(*a*) = P(*Y* | *A*=*a*) is the proportion of the population who are seropositive at age a and *F*′(*a*) is the derivative of *F*(*a*) with respect to *a*. Key assumptions include stationarity/homogeneity (i.e., no intervention or cohort effects) and that there is no seroreversion [33]. There was no evidence for large changes in transmission during the studies, even due to intervention (Supplementary Information File 1). We know for many enteric pathogens children in the Kenya cohort did serorevert (e.g., Figure 6A); when assumption is violated, estimates provide a lower-bound of a pathogen’s force of infection. We considered three different estimation approaches for force of infection from age-structured seroprevalence.

#### Exponential model (SIR model)

The simplest catalytic model, a susceptible-infected-recovered (SIR) model, assumed a constant force of infection over different ages, *λ*(*a*) = *λ* and no seroreversion [80]. In the survival analysis context, this is equivalent to assuming a constant hazard, which can be estimated with an exponential survival model. We modeled the probability of being seropositive conditional on age with a generalized linear model fit with maximum likelihood that assumed a binomial error structure and complementary log-log link [37]. We estimated average force of infection

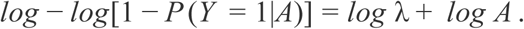

#### Reversible catalytic model (SIS model)

For some infectious diseases, like malaria, reversible catalytic models have been proposed to estimate force of infection from an age-seroprevalence curve while accounting for antibody waning with time since infection [34,81]. The model assumes a constant rate of seroconversion, λ, but extends the SIR model by also assuming a constant seroreversion rate, ρ, equivalent to a susceptible-infected-susceptible (SIS) model. The probability of a child being seropositive conditional on age is then modeled as a function of these two additional parameters:

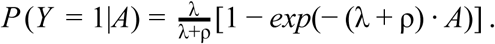

As we show in Supplementary Information File 6, age-dependent seroprevalence curves technically contain no information about seroreversion (ρ). Therefore, we fit the model with maximum likelihood assuming a binomial error structure and also assuming a fixed seroreversion rate equal to the parameters estimated separately in the longitudinal analysis (Figure 6A). As an internal validity check, we confirmed that force of infection estimates from the SIS model matched those from the SIR model for ETEC and *Campylobacter*, pathogens which had seroreversion rates that approached 0.

#### Semiparametric spline model

We fit a model that allowed force of infection to vary flexibly by age using cubic splines in a generalized additive model [71]. Let η[*P* (*Y* = 1|*A*)] = *logit P* (*Y* = 1|*A*) = *g*(*A*) for an arbitrary function *g*(·), which we fit with cubic splines that had smoothing parameters chosen through cross-validation. The age-specific prevalence predicted from the model is:

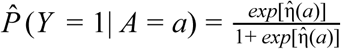 and the age-specific force of infection is:

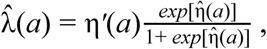

where η′(*a*) is the first derivative of the linear predictor from the logit model [33]. We estimated η′(*a*) and its standard error using finite differences from spline model predictions [71,72]. To estimate average force of infection from the model, comparable to other methods used, for each pathogen we estimated the marginal average force of infection over the empirical age distribution in the cohort:

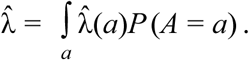

We estimated approximate 95% confidence intervals for the average force of infection by simulation with a parametric bootstrap that was based on the posterior distribution of model parameters [75,82]. Supplementary Information File 6 provides additional details, including age-specific estimates of force of infection for each pathogen.

## Data availability and replication files

Analyses were conducted in R version 3.5.0. Data and computational notebooks used to complete the analyses are available through GitHub and the Open Science Framework (osf.io/r4av7).

## Acknowledgements

The authors are grateful to Drs. Ciara E. O’Reilly and Jennifer L. Murphy for oversight and field assistance with dried blood spot collection in the Kenya study, and to Katy Hamlin for assistance with the analysis of specimens in the Haiti study. We thank William Petri and Joel Herbein for the kind gift of LecA antigen.

## Supplementary Information Files

**Supplementary Information File 1.**

No effect of intervention or bead lot on enteropathogen antibody response in Kenya and Tanzania (osf.io/vdp9a).

**Supplementary Information File 2.**

Classification agreement between different seropositivity cutoff approaches (osf.io/7x6sw).

**Supplementary Information File 3.**

Joint distributions of antibody response (osf.io/wchzq).

**Supplementary Information File 4.**

Sensitivity analysis of change in IgG used to identify presumed unexposed measurements in Haiti and Kenya (osf.io/nqrg3).

**Supplementary Information File 5.**

Estimation of age-dependent means and seroprevalence using multiple approaches (osf.io/3j79a).

**Supplementary Information File 6.**

Estimation of force of infection from age-structured seroprevalence in Kenya (osf.io/nc67w).

---

**Figure 1-supplement 1.**
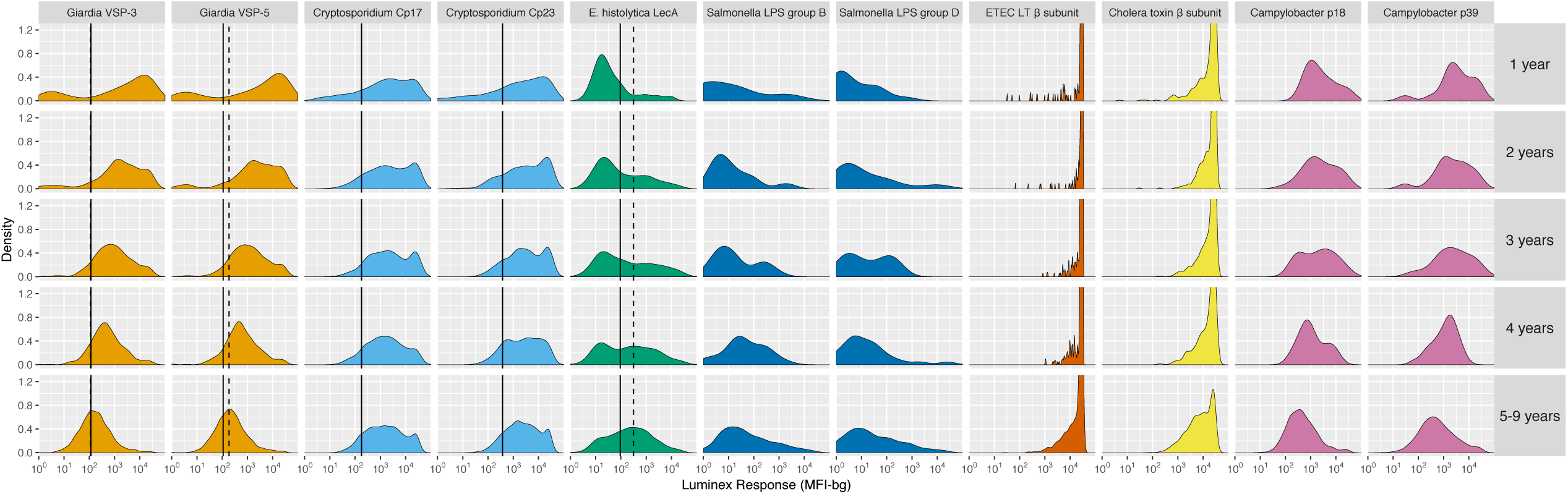
Age-stratified, IgG distributions among 4,989 children ages 1 to 9 years old in Kongwa, Tanzania, 2012–2015. IgG response measured in multiplex using median fluorescence units minus background (MFI-bg) on the Luminex platform. Vertical lines mark seropositivity cutoffs based on ROC analyses (solid) and finite Gaussian mixture models (heavy dash). Mixture models failed to converge for all antibody distributions except for *Giardia sp.* and *E. histolytica*. Created with notebook (https://osf.io/dt9zu) and data (https://osf.io/kv4d3).

**Figure 1-supplement 2.**
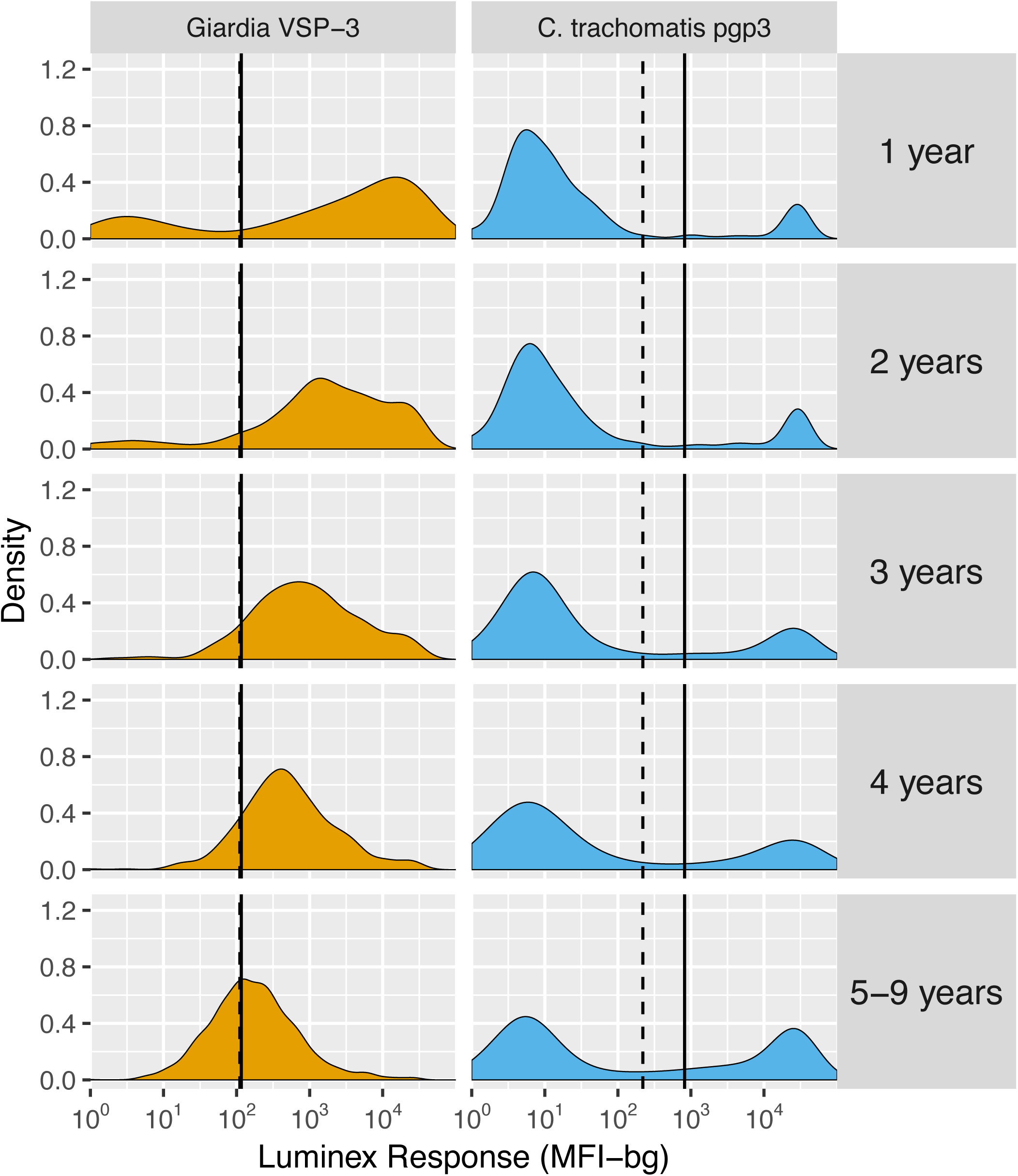
Contrasting age-dependent changes in distributions of IgG levels to *Giardia* sp. VSP-3 and *Chlamydia trachomatis* pgp3 antigens among 4,989 children ages 1 to 9 years old in Kongwa, Tanzania, 2012–2015. IgG response measured in multiplex using median fluorescence units minus background (MFI-bg) on the Luminex platform. Vertical lines mark seropositivity cutoffs based on ROC analyses (solid) and finite Gaussian mixture models (dash). Created with notebook (https://osf.io/dt9zu) and data (https://osf.io/kv4d3).

**Figure 1-supplement 3.**
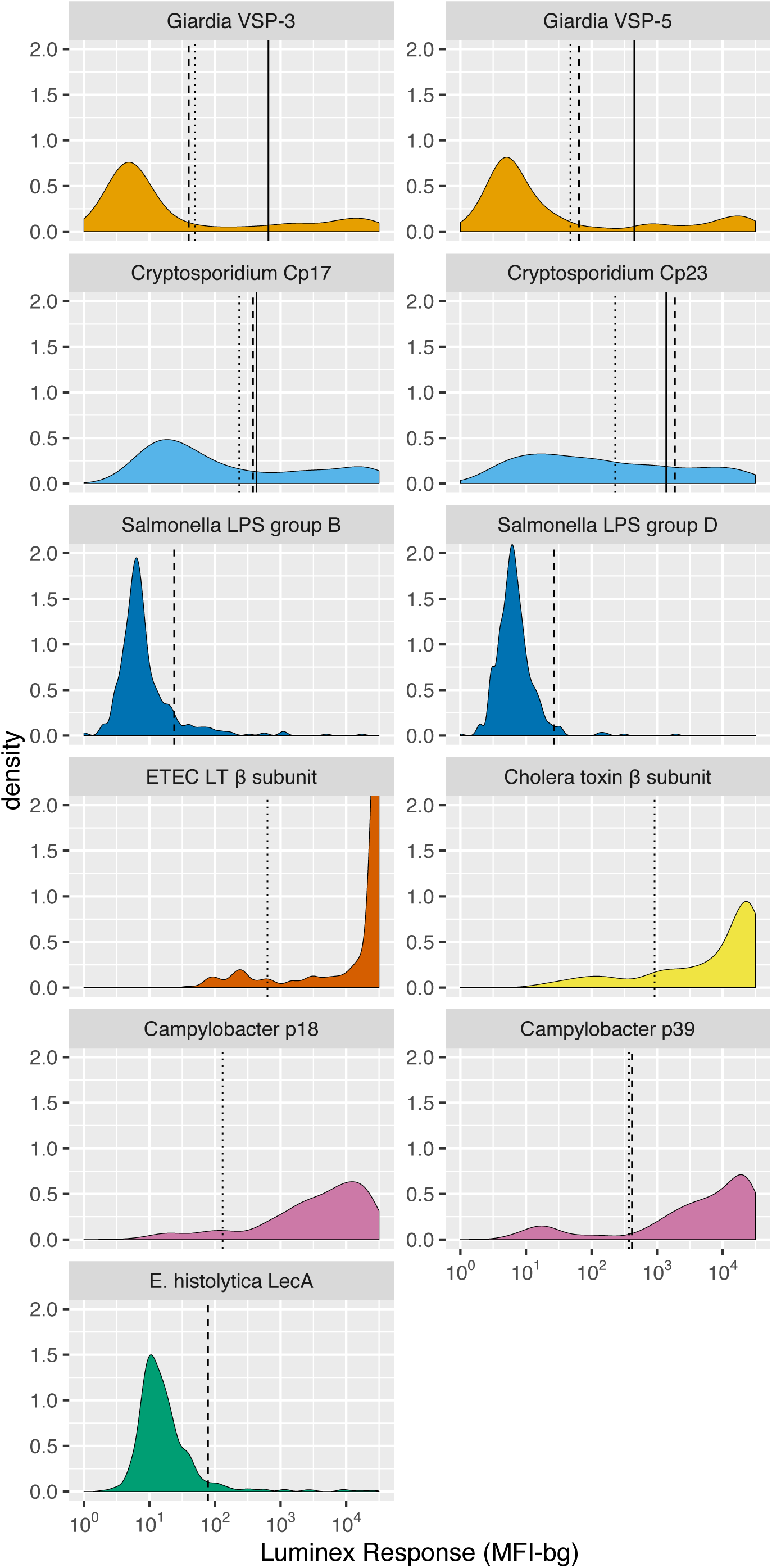
IgG distributions among children ages 4 to 17 months old in Asembo, Kenya, 2013. IgG response measured in multiplex using median fluorescence units minus background (MFI-bg) on the Luminex platform. N=439 samples from 240 children. Vertical lines mark seropositivity cutoffs based on ROC analyses (solid), finite Gaussian mixture models (heavy dash), or distribution among presumed unexposed (light dash). Mixture models failed to converge for ETEC heat labile toxin *β* subunit and *Campylobacter spp.* p18. Created with notebook (https://osf.io/456jp) and data (https://osf.io/2q7zg).

**Figure 3-supplement 1.**
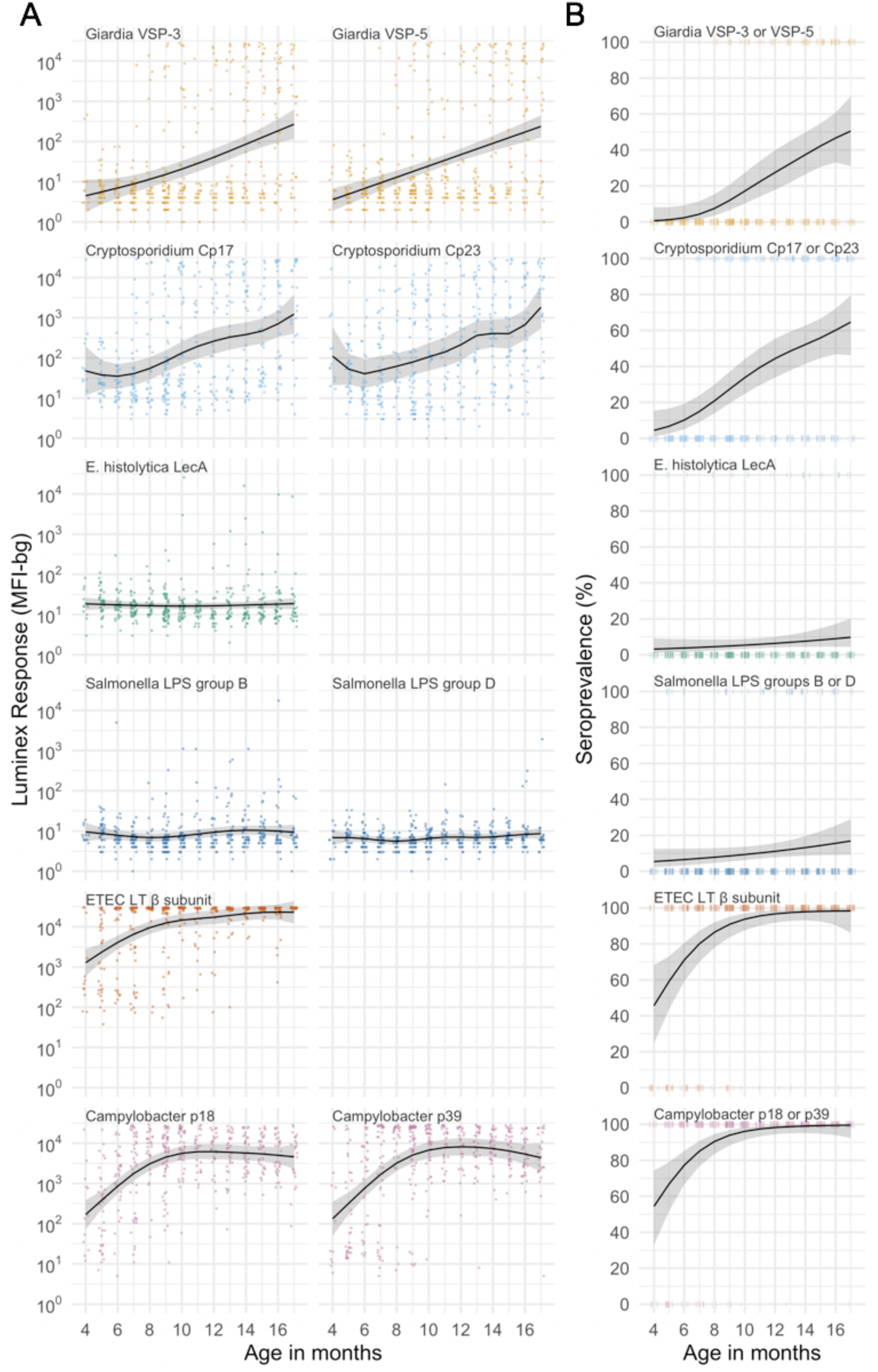
Age dependent curves for geometric means (**A**) and seroprevalence (**B**), estimated with cubic splines among children ages 4 to 17 months in Asembo, Kenya, 2013. Shaded bands are approximate, simultaneous 95% credible intervals. Ages were measured in months completed (rounded) so points in the figure are jittered. IgG response measured in multiplex using median fluorescence units minus background (MFI-bg) on the Luminex platform (N=439 measurements from 240 children). Created with notebook (https://osf.io/6c7j8) and data (https://osf.io/2q7zg).

**Figure 3-supplement 2.**
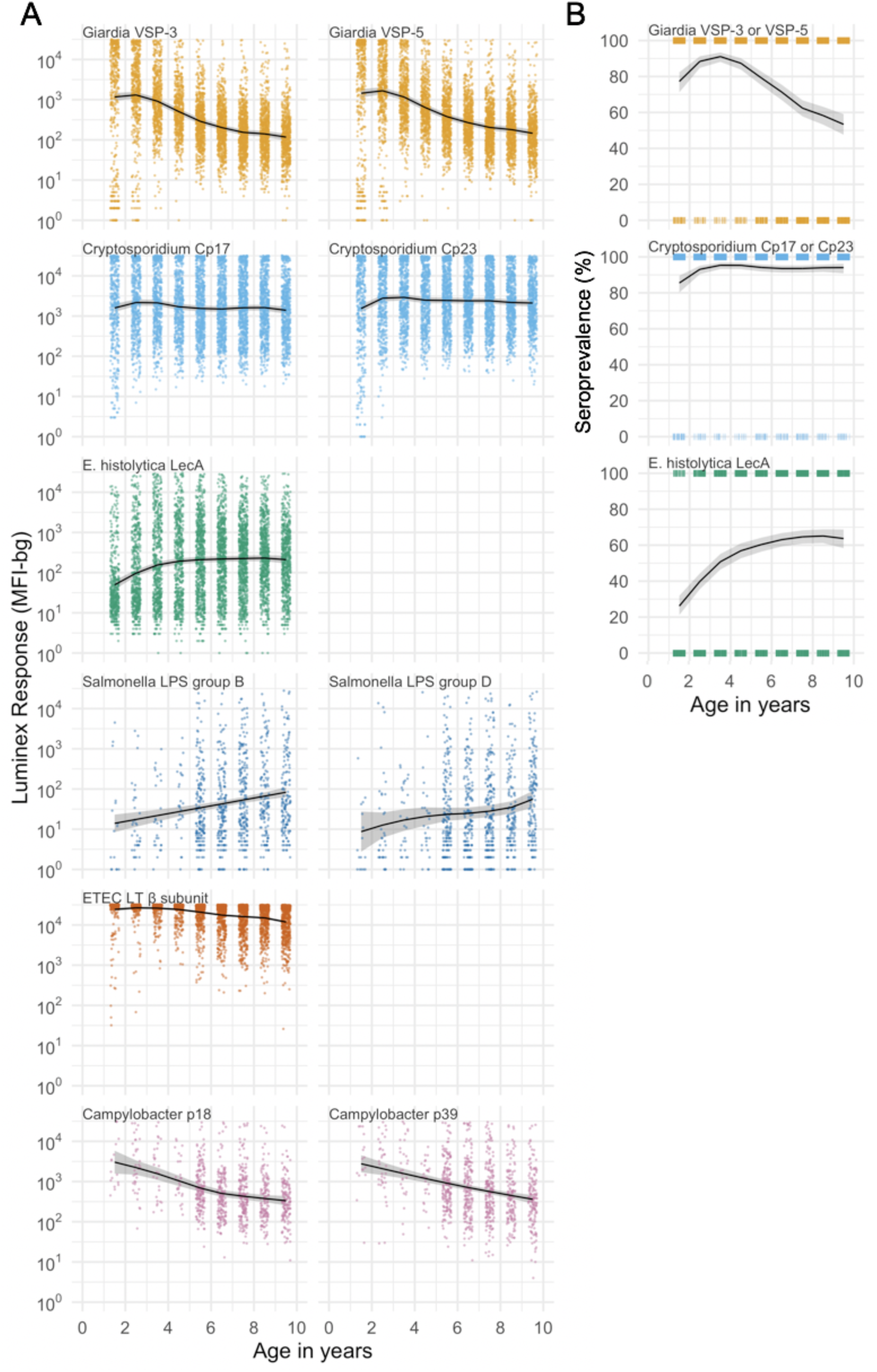
Age dependent geometric means (**A**) and seroprevalence (**B**), estimated with cubic splines among children ages 1 to 9 years in Kongwa, Tanzania, 2012-2015. Shaded bands are approximate, simultaneous 95% credible intervals. Ages were measured in years completed (rounded) so points in the figure are jittered. IgG response measured in multiplex using median fluorescence units minus background (MFI-bg) on the Luminex platform in 4,989 specimens. *Salmonella* sp. and *Campylobacter* sp. antigens were only included in the multiplex in 2012 (N=902 specimens). Seropositivity cutoffs could not be estimated for bacterial pathogens in this study, so seroprevalence curves are not shown. Created with notebook (https://osf.io/6c7j8) and data (https://osf.io/kv4d3).

**Figure 4-supplement 1.**
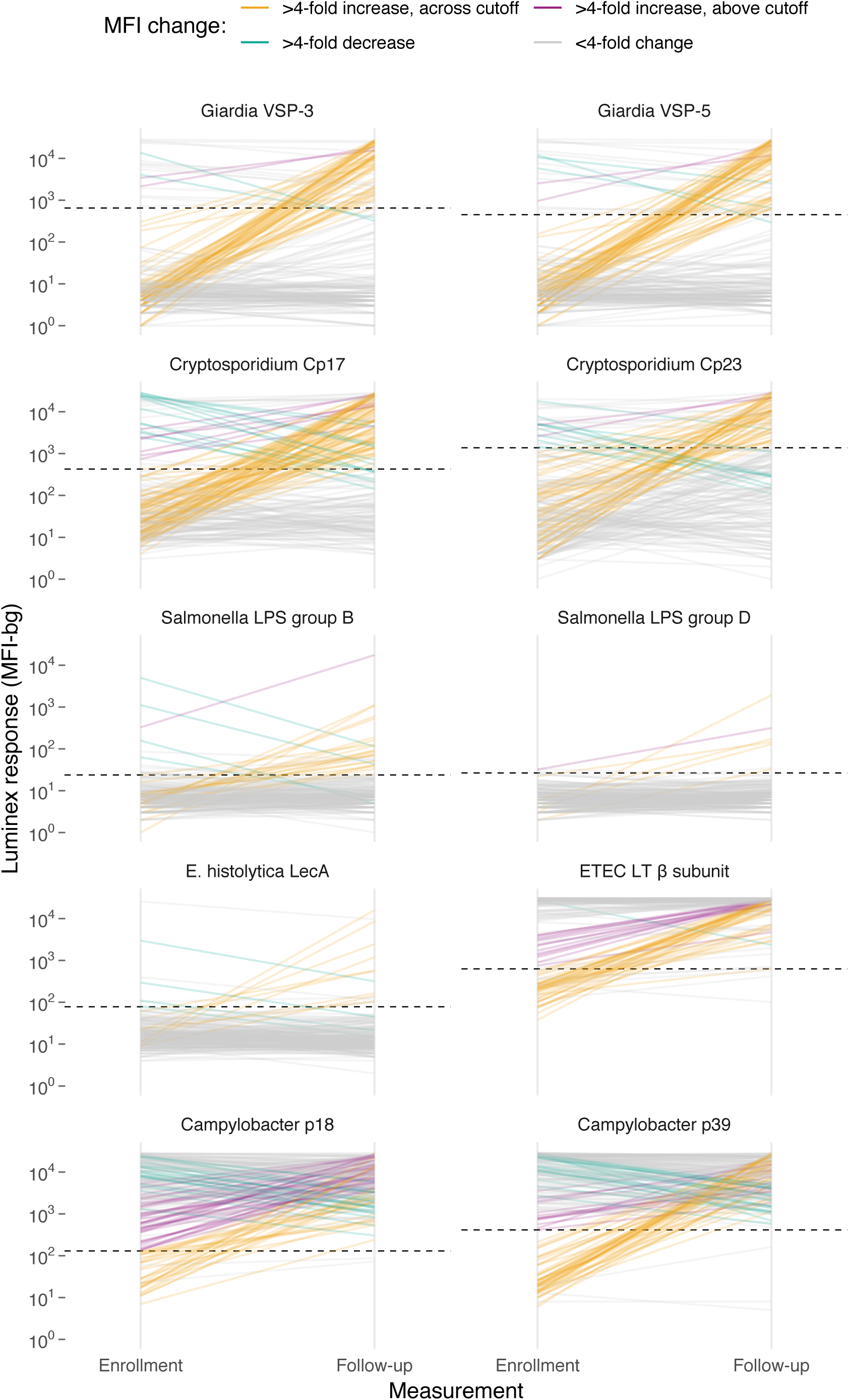
Longitudinal changes in IgG response between enrollment and followup among 199 children ages 4-17 months in Asembo, Kenya, 2013. Horizontal dashed lines mark seropositivity cutoffs for each antibody. IgG response measured in multiplex using median fluorescence units minus background (MFI-bg) on the Luminex platform. Created with notebook (https://osf.io/qhv5z), which includes additional visualizations, and data (https://osf.io/2q7zg).

